# The distress context of social calls evokes a fear response in the bat *Pipistrellus* abramus

**DOI:** 10.1101/2023.06.11.544437

**Authors:** Kazuki Yoshino-Hashizawa, Yuna Nishiuchi, Midori Hiragochi, Motoki Kihara, Kohta I Kobayasi, Shizuko Hiryu

## Abstract

Bats primarily use sound information, including echolocation, for social communication. Bats under stressful conditions, for example when confronted by a predator, will emit aggressive social calls. The presentation of aggressive social calls, including distress calls (DCs), is known to increase heart rate (HR), but how this change in HR is related to the bat’s sound perception and how this evokes behaviors such as the fear response is unknown. Herein, we show that the perception of a distress context induces freezing behavior as a fear response in bats. We found that bats responded by freezing and displayed increased HRs when they were presented with a conspecific donor bat in a distress situation evoked by gentle poking with a cotton swab. In addition, when we presented two types of auditory oddball paradigms with different probabilities of DCs and echolocation calls (ECs), the bats’ HRs increased when DCs were presented as deviant or control stimuli within standard ECs but did not increase when DCs were presented as standard stimuli. These results suggest that the situational context created by the frequency of sound presentation, rather than simply a single sound feature, induces HR increases and freezing as fear responses in bats.

**Summary statement:** We investigated the electrocardiograms of captive *Pipistrellus abramus* and found that their heart rate increased as a fear response when the bats heard sounds with a distress context.

## Introduction

Echolocating bats rely heavily on sound information as both a means of learning about their external environment and as a factor in determining their own behavior. Bats are social animals that communicate primarily through sound (Chaverri et al., 2018). The social calls of echolocating bats include aggressive calls, warning calls, mating calls, songs, and isolation calls; which one is used depends on the context (Bohn et al., 2008; Davidson and Wilkinson, 2004; Fenton, 2003; Gelfand and McCracken, 1986; Pfalzer and Kusch, 2003). For example, distress calls (DCs), a form of social call, are produced by many animals when they are in situations of extreme danger such as being caught by a predator or tangled in a catch net (Carter et al., 2015; Conover, 1994; Lingle et al., 2012; Ruiz-Monachesi and Labra, 2020), and bats express DCs when in a stressful or dangerous situation (Chaverri et al., 2018; Fenton et al., 1976). Previous studies have reported the possible roles of the DCs of bats as direct responses to predators, as warnings to kin or non-kin individuals, and as means of attracting conspecifics or heterospecifics for mobbing predators (Arnold et al., 2022; August, 1985; Carter et al., 2015; Knörnschild and Tschapka, 2012; Russ et al., 2004; Ryan et al., 1985). In addition, the acoustic structures of such vocalizations may contain information about the caller, such as body size, health status, and fear status (August and Anderson, 1987; Gadziola et al., 2012; Hechavarría et al., 2020; Jiang et al., 2017). In addition to the acoustic analysis of these social calls and research concerning their functions, it is also important to study changes in the internal states of the bats receiving social calls. For example, it has been reported that DCs evoke changes in neurotransmitters and stress hormones in the amygdala as a fear-related response in bats (Mariappan et al., 2013). The heart rate (HR) is an important parameter in the evaluation of internal states, as it can indicate tense, aggressive, or appeasement states evoked via the autonomic nervous system. A previous study reported that the magnitude and duration of elevated HRs were correlated with the level of evoked aggression in emitter bats (Gadziola et al., 2012). Some studies have used changes in HR to detect the fear level in social calls, and these studies have reported that aggressive stimuli evoked HR increases as a fear response in receiver bats (Gadziola et al., 2015; Hechavarría et al., 2020; Ma et al., 2010). Thus, the usefulness of HR changes for assessing internal states, such as emotions, has been demonstrated in studies of bats focusing on the recipients of social calls. The presentation of aggressive social calls, such as DCs, is known to increase the receiver bats’ HR, but how the HR change is based on the bat’s perception and how this leads to a given behavior (i.e., a fear response) have yet to be investigated. In particular, recent studies have shown that context consisting of several vocalization sequences, rather than a single sound stimulus, may be important in bats’ communications (Amit and Yovel, 2023). Thus, we expected that, instead of a single DC stimulation, the recognition of a distress context or situation would increase the HR as a fear response. In this study, therefore, we 1) investigated whether fear responses (freezing) and HR increases were observed in subject bats when confronted with the stimulus of a donor bat in distress emitting a DC and 2) investigated whether the acoustic context of the DCs evoked a fear response using the auditory oddball paradigm which consists of different presentation probabilities of DCs. We examined the fear response and cognitive processes in a distress context through these experiments.

## Materials and Methods

### Study species and ethical status

The study species was the Japanese house bat *Pipistrellus abramus*, an insectivore and aerial hunter in the order Vespertilionidae. For echolocation, this species uses frequency-modulated (FM) pulses that sweep downward from initial frequencies around 80 kHz to terminal frequencies (TF) around 40 kHz (as recorded in the field: Fujioka et al., 2011; Hiryu et al., 2008). Auditory sensitivity exhibits a broad U-shape ranging from approximately 4 kHz to 80 kHz and a low threshold between 20 and 50 kHz, with the highest sensitivity in the 35–50 kHz frequency range (Boku et al., 2015; Goto et al., 2010).

Subjects were captured from a colony roosting near the campus of Doshisha University in Kyotanabe, Japan, or were birthed in our laboratory from captive pregnant females. The bats were kept in plastic cages (25 cm×17 cm×17 cm) and allowed free access to mealworms and water. Eighteen adult bats (8 males and 10 females) were used in this experiment. All experiments complied with the Principles of Animal Care (publication no. 86-23 [revised 1985] of the National Institutes of Health) and all Japanese laws. All procedures were approved by the Animal Experiment Committee at Doshisha University.

### Experiment I: Behavioral response to a donor bat stimulus

We recorded sounds in the experimental environment and the behavioral responses of subject bats when confronted with a donor bat in a state of distress. Six bats (four females and two males) were used in this experiment. The experiment was carried out in a cylindrical arena (70 cm in diameter, 50 cm in height) surrounded by a metal net wall in a soundproof chamber (1.4 m×1.4 m×2.1 m) with red light (G-82H(R), ELPA, Japan) that prevented the animals from relying on visual information (Hope and Bhatnagar, 1979). A video camera (DMK33UX273, The Imaging Source, Germany, frame rate: 10 fps) was set 70 cm above the center of the arena to observe the entire area. Acoustic signals were recorded with a microphone (Anabat SD2 CF Bat Detector, Titley Scientific, Australia) placed on the arena floor. The sound recordings were conducted through a data acquisition system (BNC-2110 / PXIe-6358 / PXIe-1073, NI, USA) and a custom-made program in Labview 2016 (NI, USA, sampling rate: 500 kHz/channel).

Three adult male bats were used as stimulus donors, all of which were kept in separate rearing cages from the time of capture. A stimulus donor bat was placed in a metal grid cage (17 cm × 17 cm × 17 cm) with an open top. We set three conditions of stimuli in this experiment: “Distressed bat (DB)”, “Non-distressed bat (NDB)”, and “Control” (no bat in the chamber) conditions. In the DB condition, the experimenter gently poked the donor bat with a cotton swab to evoke a distress call as in previous studies (Gadziola et al., 2012). For the NDB condition, the donor bat was able to move freely within the cage without a stress stimulus. For the control, only the cage (without a donor bat) was present in the arena.

To prevent echolocation until the recording began, the experimenter gently placed a subject bat covered with a plastic cup (5 cm in diameter and 3 cm high) at the starting point, 30 cm away from the center of the arena. Then, the cage with the stimulus donor bat (no bat in the control condition) for each condition was placed in the arena about 20 cm away from the subject bat. As soon as recording began, a rope attached to the top of the cup was pulled to release the cover. Each condition was presented for three minutes (one trial), and we conducted a total of 31 trials for six subject bats (DB: 10 trials, NDB: 10 trials, Control: 11 trials; Table S1).

The response behaviors of the subject bats were classified as “fly”, “crawl”, or “stay”. The reaction time from the start of the stimulus (the time when the cup over the subject bat was removed) until the subject “flew” or “crawled” was also recorded (Figure S1). Sound data were analyzed by a custom-made program in MATLAB (R2019b, MathWorks, USA). One hundred calls that were randomly selected from recorded calls 6 dB p-p greater than the noise level were manually classified as echolocation calls (EC) or distress calls (DC). Note that the DCs were emitted only by the donor bat, while the ECs from the donor bat were not distinguished from those of the subject bat. The DCs were further classified into three types as follows: “DCFM” (down FM pattern with lower TF than echolocation calls and a duration shorter than 10 ms), “DCNB” (noise burst lasting more than 10 ms), or “DCO” (calls that were not classified as either DCFM or DCNB). The initial frequency (IF) and the TF of DCFM were analyzed based on a spectrogram (2048 point windows, −25 dB threshold from the peak power) to check the distribution as the stimuli for experiment III.

### Experiment II: ECG measurement for behavioral stimuli

In this experiment, we measured the electrocardiogram (ECG) as a physiological parameter to examine the internal states of the subject bats in real time via autonomic nervous system functions. Five adult bats (two females and three males) were used as subjects in this experiment. Subject bats were anesthetized with 2–4% isoflurane (Pfizer, USA) during surgery, and 2% xylocaine jelly (Sandoz, Switzerland) was applied to the surface of the skin after the scalp fur was shaved. Then, the skin and muscles over the cranium were retracted. The ECG wires were directed to the electrode socket and fixed with dental cement (Super-Bond C&B, Sun Medical Company, Japan / UNIFAST III, GC, Japan) on the cranium. These procedures referred to our previous studies (Boku et al., 2015, Furuyama et al., 2018). The ECG electrodes (silver wires) were implanted in the subject bat in three different configurations (Lead II) as follows: left leg (as positive), right thumb (as negative), and right leg (as a reference) and fixed under the skin. Lead II is one of the highest signal-to-noise ratio measurements for ECG in echolocating bats (Mihova and Hechavarría, 2016).

This experiment was carried out in the same soundproof chamber using the same behavioral recording system as in experiment I. Subject bats participated in the experiment after at least one week of recovery following the surgery. On the day of the experiment, the subject bat was immobilized with a soft sponge and placed on the arena floor approximately 20 cm away from the stimulus donor cage. The electrode socket of the subject was connected to an ECG measurement system (C3410 / C3314 / C3216 / C3100, Intan Technologies, USA). Before the recording, we tested whether the electric impedance of all ECG electrodes was under tens of kiloohms before the measurement to check that the electrodes were functioning. The ECG was recorded by RHX software (Intan Technologies, USA) using 30 kHz sampling and was synchronized to video recordings by a pull-down trigger signal.

Five adult bats (three females and two males) were used as stimulus donors. Four of the donor bats were kept in separate rearing cages from the time of capture of the subject bats. One subject–donor pair was kept in the same cage from the time of capture. All pairs were of the same sex to avoid sexual interaction (the details are listed in Table S2). We set the DB and NDB conditions the same as in experiment I, but two control conditions were set: one in which only the cage without a stimulus donor bat was presented (Control I) and the other in which no cage was presented (Control II). These four conditions were presented randomly, and each condition was presented to the subject bat for at least one minute (one trial). The ECG recordings were stopped depending on the poor signal-to-noise ECG condition of the subject or the fatigued appearance of the donor bats so that the recording time ranged from 8 to 21 minutes.

### Experiment III: ECG measurement for acoustic stimuli

To avoid confounding stimuli, such as odor from the donor bat or movements of the experimenter, and to measure responses based on acoustic information, the ECG was recorded while pre-recorded acoustic stimuli (calls emitted by a donor bat) were presented from a loudspeaker instead of by a donor bat. Note that the acoustic parameters of the DC stimuli in this experiment were similar to the average level of the DCFM in experiment I, as shown in Fig. S2.

The experiment was carried out in the same soundproof chamber as in experiment I, and the surgical procedure was the same as in experiment II. The loudspeaker (PT-R7III, Pioneer, Japan) was installed on the floor of the arena in the chamber and connected to an amplifier (STR-DH190, Sony, Japan) and D/A converted via an audio interface (Rubix44, Roland, Japan, sampling rate: 192 kHz). The stimulus calls were pre-recorded by a sound acquisition system (CM16/CMPA / UltraSoundGate 116Hme, Avisoft-Bioacoustics, Germany) from a male donor bat while the experimenter held the bat in his hand to elicit ECs and DCs as in previous studies (Carter et al., 2015; Hörmann et al., 2021; Jiang et al., 2017; Russ et al., 2004). Here, the experimenter handled the donor bat, maintaining a constant distance between the microphone and the bat, as the loudspeaker was fixed as the presentation system. Each recorded sound sequence was trimmed into a 3-second echolocation call period and a 1.5-second distress call period. We selected the DCFM type as the distress stimulus in this experiment because a previous study reported that the DCFM type increased heart rate (HR) by more than the DCNB type (Gadziola et al., 2015). Then, successive vocalizations were normalized for amplitude and adjusted to a length of three seconds by repeating the distress call word cutoff at 1.5 seconds twice. These stimuli were presented at 100 dB p-p SPL to the subject bat at about 10 cm from the center of the loudspeaker. Here, we adopted the flip-flop auditory oddball paradigm (Näätänen et al., 1978; Ulanovsky et al., 2003) to examine changes in HR induced by the acoustic stimuli rather than unexpectedness.

The distress calls and the echolocation calls were used as stimuli in this paradigm. This paradigm comprised standard stimuli presented at a 90% probability of occurrence and deviant stimuli presented at a 10% probability of occurrence. At least three standard stimuli were inserted after each deviant stimulus to maintain the deviance effect. The deviant stimuli and standard stimuli occurred in a random series once after every silence interval. The interval was set at 30 s to return to the pre-stimulation HR level, referencing an HR study for an acoustic task in birds (Ikebuchi et al., 2003). At the end of the trial, the same stimulus as the deviant was played 10 times continuously, and this was used as the control session. Therefore, one trial contained a total of at least 110 stimuli (90 standard, 10 deviant, and 10 control stimuli). Note that the number of presentations of the standard stimuli was increased due to the forced insertion of the standard stimuli after the deviant stimuli. We set two types of oddball paradigms by flipping the standard and deviant stimuli. We conducted 11 trials of the EC- standard oddball paradigm using six bats (four females and two male subjects) and five trials of the DC-standard oddball paradigm with four bats (two females and two male subjects from the default oddball paradigm participated). For each individual bat, measurements lasted approximately one hour.

### ECG signal analysis

The ECG signal processing was conducted using a custom-made MATLAB script (R2019b, MathWorks, USA). The ECG was obtained from the potential difference of two electrodes between the positive and negative terminals attached to the subjects. The ECG was bandpass filtered (40 to 150 Hz) to detect the peak of the R waves and to eliminate outliers based on the amplitude distribution (a range of 5% to 95% was retained). The instantaneous HR was calculated from the inverse of the valid R wave interval of the ECG. To analyze the effect of the stimuli, the HR was binned by 1 s before/after the onset of the stimuli via averaging. The binned HR was analyzed using a −10 to 90 s period in experiment II and a −10 to 30 s period in experiment III that was fitted to each interstimulus interval (where the stimulus onset was set to 0 sec). Finally, the HR was normalized for the average of each −10 to 0 s period for the onset to evaluate values that considered individuality and temporary activeness and thus the relative effect of the stimuli.

### Statistical analysis

All statistical analyses were performed using a self-written MATLAB script (R2023a, MathWorks, USA). To test for significant deviation from a normal distribution, we used the MATLAB “lillietest” function. Based on the results, we used post hoc non-parametric tests comparing average ranks (Matlab functions “kruskalwallis” and “multcompare,” with the Bonferroni correction) or parametric tests for the relative HR bins after the stimulation period with the stimulation group in experiments II and III. In our case, we used only non-parametric tests because all data deviated from a normal distribution.

## Results

### Experiment I: Behavioral response to stimulus donor bats

In 10 trials using the DB condition, a total of 11,187 calls were recorded from three male donor bats. Figure 1A shows example vocalization sequences of two donor bats; the vocalizations have complex temporal patterns, including DCFM and DCNB types. In all 10 trials with the DB condition, DCNB had the highest vocalization rate (Fig. 1B). The average duration of EC in *P. abramus* was 1.12 ± 0.33 ms (mean ± SD), whereas the duration of DCNB was the longest (71.77 ± 45.1 ms), at times exceeding 100 ms (Fig. 1C).

**Fig. 1.**
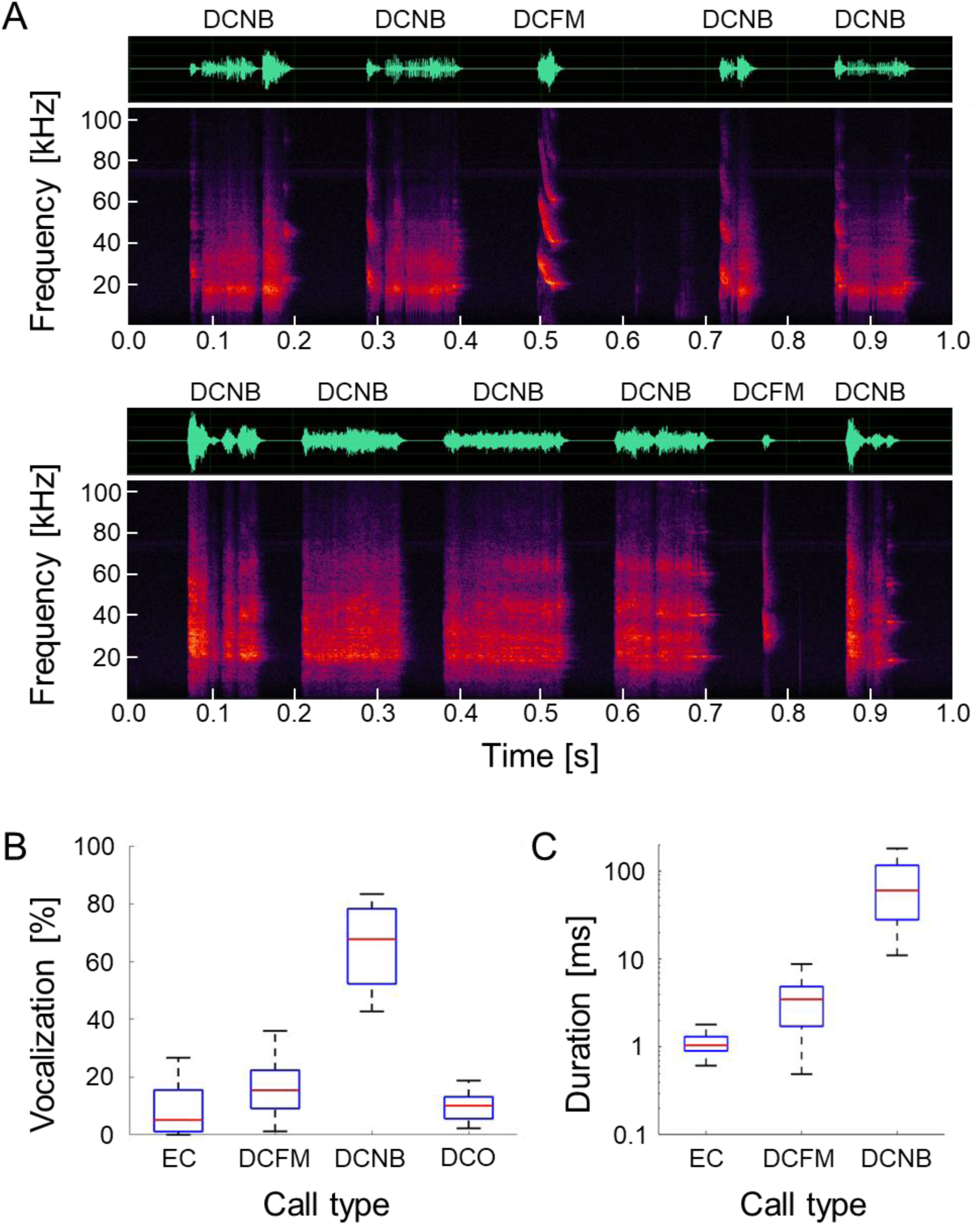
Vocalizations of the donor *P. abramus* in the distress situation. (A) Example of a series of distress vocalizations from two different individuals. Each panel shows a time waveform (top) and a spectrogram (bottom). (B) Box plots of vocalization rate versus call type. (C) Box plots of duration versus call type. Here, DCO was eliminated due to a lack of classification rules. Each box plot shows the median (red line), quartiles (the bottom and top edges of the boxes), and the whiskers (the minimum and maximum) (the top and the bottom tips of the vertical lines).

Figure 2 shows the percentages of the behavioral responses of the subject bats in each condition (the result of each trial is shown in detail in Table S1). In the DB condition, subjects were most likely to respond with “stay” (60.0%) and least likely to respond with “fly” (Fig. 2). In the NDB condition, in contrast, “fly” was the dominant response at 50.0%. In the control condition, the percentage of “stay” was 27.3% and those of “crawl” and “fly” were both 36.4%.

**Fig. 2.**
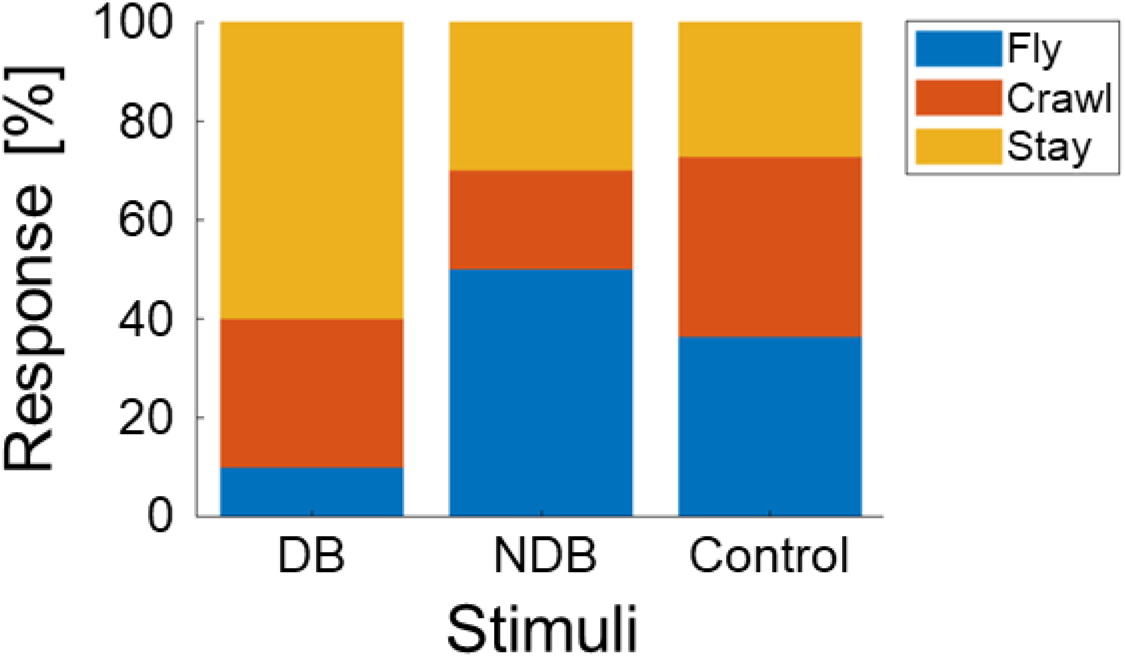
Distributions of behavioral responses in the experiment. The DB and NDB stimuli each show 10 responses, and the control shows 11 responses.

### Experiment II: ECG measurement for behavioral stimuli

We conducted 60 trials (DB: 15 trials, NDB: 21 trials, Control I: 9 trials, Control II: 15 trials; see Table S2 for details, including information on donor–subject bat combinations). Recorded ECGs had stable S/N ratios with no clips (approximately ± 0.1 mV range) in all trials, as shown in Fig. 3A. The ECGs included clear R, S, and T waves, as in other species (Currie, 2018). The HR ranged from approximately 200 to 600 bpm through all measurements. Figure 3B shows an example comparing the change in relative HR during 90 seconds after stimulus presentation in one individual in the DB and NDB conditions. The relative HR increased by a maximum of about 40% after the onset of the DB condition despite being kept after the onset of the NDB condition. The HR average from 0 to 90 s after the onset of the DB condition was significantly higher than in the NDB condition (multiple comparison test with the Bonferroni correction, *p* < 0. 001) and control II condition (*p* < 0.05) (Fig. 3C). The DB condition also showed a marginally significant increase compared to the control I condition (*p* < 0.1). There were no significant differences between NDB and the two control conditions (*p* > 0.1, rest pairs).

**Fig. 3.**
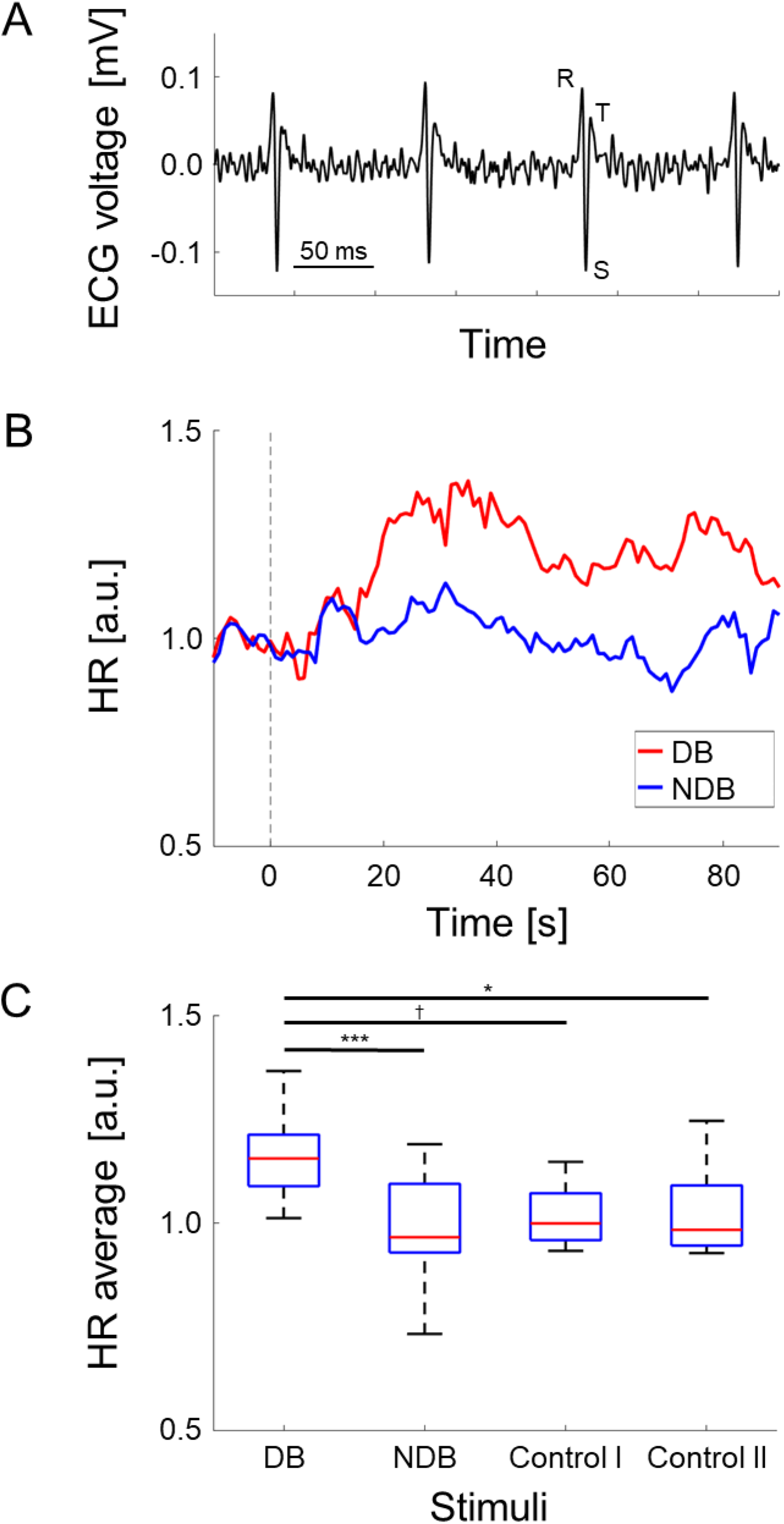
Result of ECG analysis in Experiment II. (A) Recorded ECG applied 40 to 150 Hz BPF. The R, S, and T waves are clearly visible in each cardiac beat. (B) Example of averaged relative HR responses to the DB and NDB stimuli in one subject. The vertical gray dotted line at 0 s shows the onset of stimulation. (C) Distribution of average relative HR from 0 to 90 s after the onset of stimulation. Each box plot shows the median (red line), quartiles (the bottom and top edges of the boxes), and the whiskers (the minimum and maximum) (the top and the bottom tips of the vertical lines). Statistical significance is indicated by the symbols †: P < 0.1, *: P < 0.05, ***: P < 0.001.

### Experiment III: ECG measurement for acoustic stimuli

We recorded 1255 valid responses from six bats in 11 trials using the EC-standard paradigm and 568 valid responses in five DC-standard trials. In the EC-standard paradigm, the deviant DCs caused an increase in relative HR after the onset of the stimuli (Fig. 4A). The HRs averaged from 0 to 30 s after the onset of the deviant stimuli (DCs) were significantly higher than those with the standard stimulus (EC) responses (standard vs. deviant: *p* < 0.0001; standard vs. control: *p* < 0.05; deviant vs. control: *p* > 0.1, comparisons after Bonferroni correction) (Fig. 4B). For the DC-standard paradigm, HR increases were unclear regarding the stimuli (Fig. 4C). There were no significant differences in average HR after the onset of the stimuli (standard vs. deviant: *p* > 0.1; standard vs. control: *p* > 0.1; deviant vs. control: *p* > 0.1, comparisons after Bonferroni correction) (Fig. 4D).

**Fig. 4.**
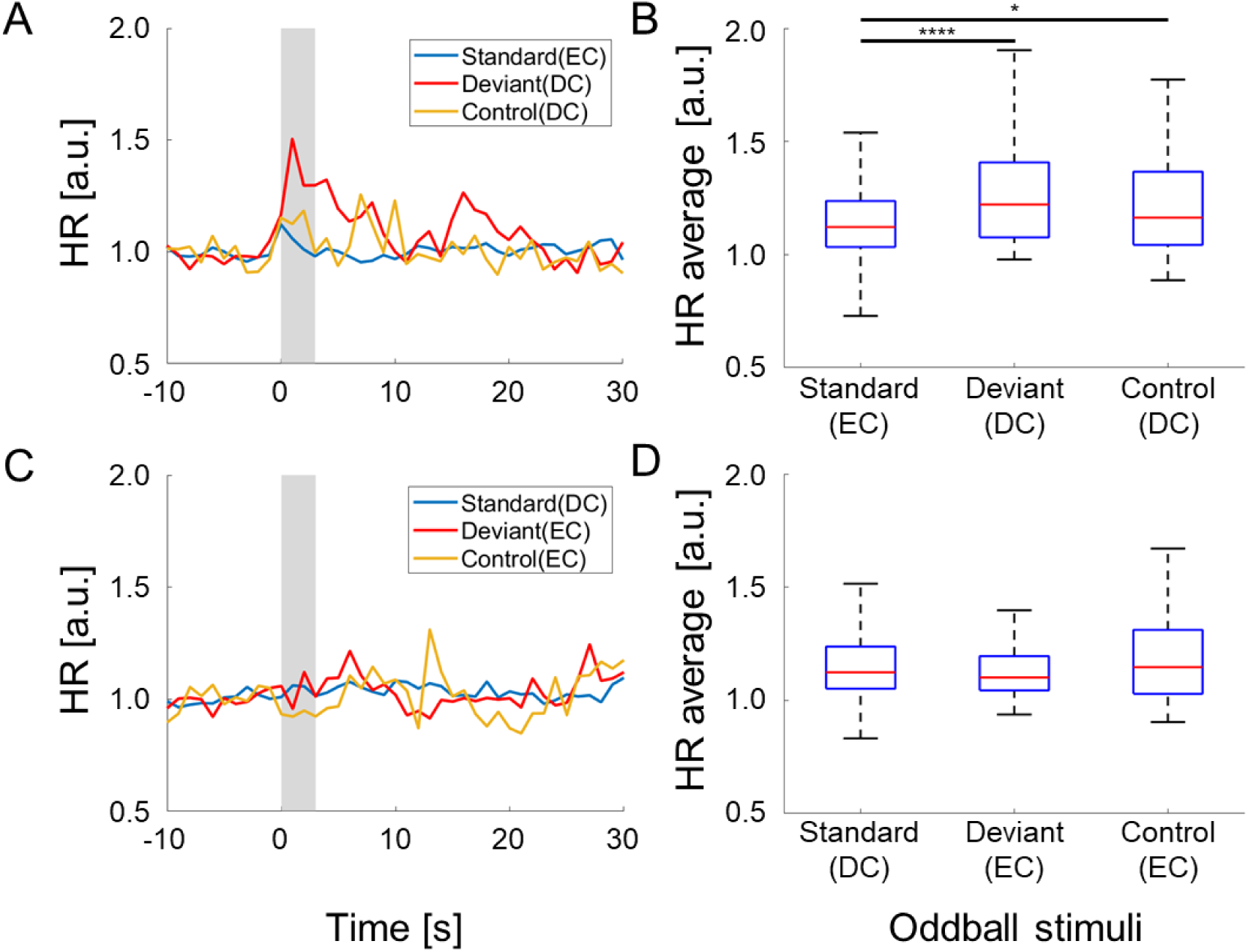
Result of ECG analysis in the oddball paradigm (experiment III). The DC-standard paradigm is shown at the top (A, B), and the EC-standard paradigm is at the bottom (C, D). Left (A, C) shows the average relative HR response from −10 to 30 s. Here, 0 s indicates the onset of stimulation, and the gray area from 0 to 3 s indicates the acoustic stimulation period. Right (B, D) shows the distribution of average relative HR from 0 to 30 s after stimulus onset. Each box plot shows the median (red line), quartiles (the bottom and top edges of the boxes), and the whiskers (the minimum and maximum) (the top and the bottom tips of the vertical lines). Statistical significance is indicated by the symbols *: P < 0.05, ****: P < 0.0001.

## Discussion

### Cognition and a distress situation

In this study, we investigated behavioral and HR responses to distress vocalizations in the Japanese house bat *P. abramus*. Using this approach, we tested the hypothesis that an HR increase in subject bats (DC receiver) is caused by the recognition of a distress situation. According to our first result, “stay” was the dominant response in the DB condition, in contrast to the majority of responses involving moving freely in the NDB and control conditions (Fig. 2). In other words, the distress context may have induced an increase in the freezing behavior as a fear response in the DC receiver. The freezing behavior has been shown to have led to a rapid increase in HR and evoked the dominance of the parasympathetic nervous system in rodents (Liu et al., 2013; Liu et al., 2014; Livermore et al., 2021). The HR increase shown in Fig. 3B may reflect the freezing behavior as a fear response. Hence, our results support the assumption that the freezing behavior was evoked by the recognition of a distress context in a conspecific by the receivers. This suggestion does not conflict with previous physiological studies related to distress context in bats (Gadziola et al., 2012; Gadziola et al., 2015; Hechavarría et al., 2020; Ma et al., 2010; Mariappan et al., 2013). However, we could not eliminate other factors such as human actions and olfactory cues. The results of the third experiment (Fig. 3A and 3B) support the assumption that the vocalization provided distress information to the receiver because the HR increased after the deviant DC stimulation within the presentation of the standard ECs. However, the DCs did not lead to HR increases during the DC-standard oddball paradigm (Fig. 3C and 3D). In other words, even with the same sound stimulus, HR responses differed depending on the type of deviation. Incidentally, we did not focus on sex or age differences in these responses because the sample size was not large enough to be statistically comparable. Future experiments will examine whether these factors elicit HR responses as different internal states. In summary, these results suggest the possibility that *P. abramus* may detect a distress situation from the vocalization context of conspecifics.

### Vocalization with distress situation in Japanese pipistrelle bats

DCs in Japanese pipistrelle bats, *Pipistrellus abramus*, had lower TF and/or longer duration than ECs (Fig. 1C), a trend similar to those of other bat species, including *Pipistrellus pipistrellus*, *Molossus molossus,* and *Carollia perspicillata* (Carter et al., 2015; Fenton et al., 1976; Gadziola et al., 2012; Hechavarría et al., 2016; Ma et al., 2006; Pfalzer and Kusch, 2003; Russ et al., 1998; Russ et al., 2004). The DCNB was categorized as a high-aggression type according to the categorization of the social calls in previous studies based on the acoustic structure patterns (Gadziola et al., 2012; Gadziola et al., 2015). In particular, the DCNB had the highest vocalization rate in our experiment. Therefore, our experimental design generally provided a high-distress situation for the subject bats in experiments I and II. In contrast, the DCFM was categorized as a low-aggression type, but this evoked greater HR increases than high-aggression calls, such as the DCNB in the receiver bats (Gadziola et al., 2015). In our study, the DCFM, which had the second-largest vocalization ratio in the distress stimulations, increased the relative HR by a maximum of 40% in the DB condition, the same as in a previous study (Gadziola et al., 2015). Although we did not compare the heart rate increase with DCNB in our present experiments, the distress context of the DCFM could evoke HR increases in *P. abramus*. The difference in HR increases between DCNB and DCFM may be related to whether the respective vocalizations of social calls are directed toward predators or to a conspecific. Further investigation is needed, but it is clear that changes in the HR are a good indicator for reading changes in internal states induced by the social calls of bats. It would be particularly interesting to investigate the HRs of wild Japanese house bats in distress situations.

The DCs could have multiple roles for conspecifics, predators, and other species (Conover, 1994; Klump and Shalter, 2010). In addition, the acoustic characteristics of the DCs in bats may be heterospecific (Arnold et al., 2022; Carter et al., 2015; Chaverri et al., 2018; Russ et al., 2004). In our results, *P. abramus* also showed similar acoustic characteristics as in other species for the DCs, as shown in Fig.1A. Therefore, there is a possibility that *P. abramus* could use information from the DCs as intraspecific and heterospecific social calls. Future experiments will test whether these DCs elicit similar HR increases in heterospecifics and whether this relies on social relationships between sender and receiver.

### ECGs of Japanese pipistrellus bats

The ECGs had clear R, S, and T waves (Fig. 3A) and were similar to those from previous studies in other bats (Currie, 2018; Mihova and Hechavarría, 2016). The P, Q, and U waves were unclear, but this is not a surprising result. The P and U waves could be clarified via averaging as they generally are of small amplitude, normally smaller than the T wave. However, the Q waves of bats remain unclear, as in rodents (Farraj et al., 2011; Kaese and Verheule, 2012; Sambhi and White, 1960). Despite such differences, there are essential similarities between bats, rodents, and humans as mammals (Konopelski and Ufnal, 2016). In this study, we focused only on HR, but other ECG parameters may contain information about the bats’ internal state. For example, cardiac arrhythmias are quite interesting, but the details of their relationship with internal states are still unclear and are extremely underreported in bats (Currie, 2018). In the future, we would like to investigate the relationship between the electrocardiogram and the internal state in more detail.

## Acknowledgments

We are grateful to Yasufumi YAMADA, Kazuma HASE, Yuta TAMAI, Yu TESHIMA, Kazuki SHIN’YA, Hidekazu NAGAMURA for their advice in recordings or writing paper.

## Competing Interests

The authors declare no conflict of interest.

## Funding

This work was supported by the Japan Society for the Promotion of Science (JSPS) KAKENHI Grant [Numbers JP 21H05295, 22H01503 to SH, 23KJ2080 to KYH], the Japan Science and Technology Agency (JST) SPRING Grant [Number JPMJST2129 to KYH], the JST establishment of university fellowships towards the creation of science technology innovation Grant [Number JPMJFS2145 to KYH] and the Japan Science Society (JSS) Sasakawa Scientific Research Grant [Number 2021-5028 to KYH].

## Data availability

Data supporting the findings of this study are available from the corresponding author upon reasonable request.

---

**Fig. S1.**
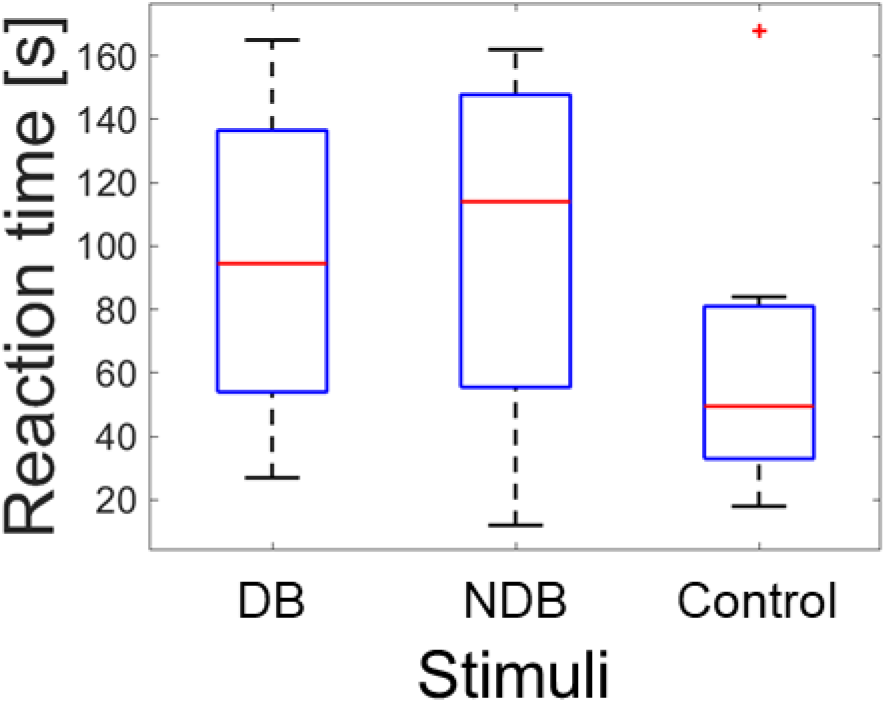
Box plots of the reaction time from experiment I. Each box plot shows the median (red line), quartiles (the bottom and top edges of the boxes), and the whiskers (the minimum and maximum) (the top and the bottom tips of the vertical lines). The red cross mark indicates an outlier (outside of the whiskers).

**Fig. S2.**
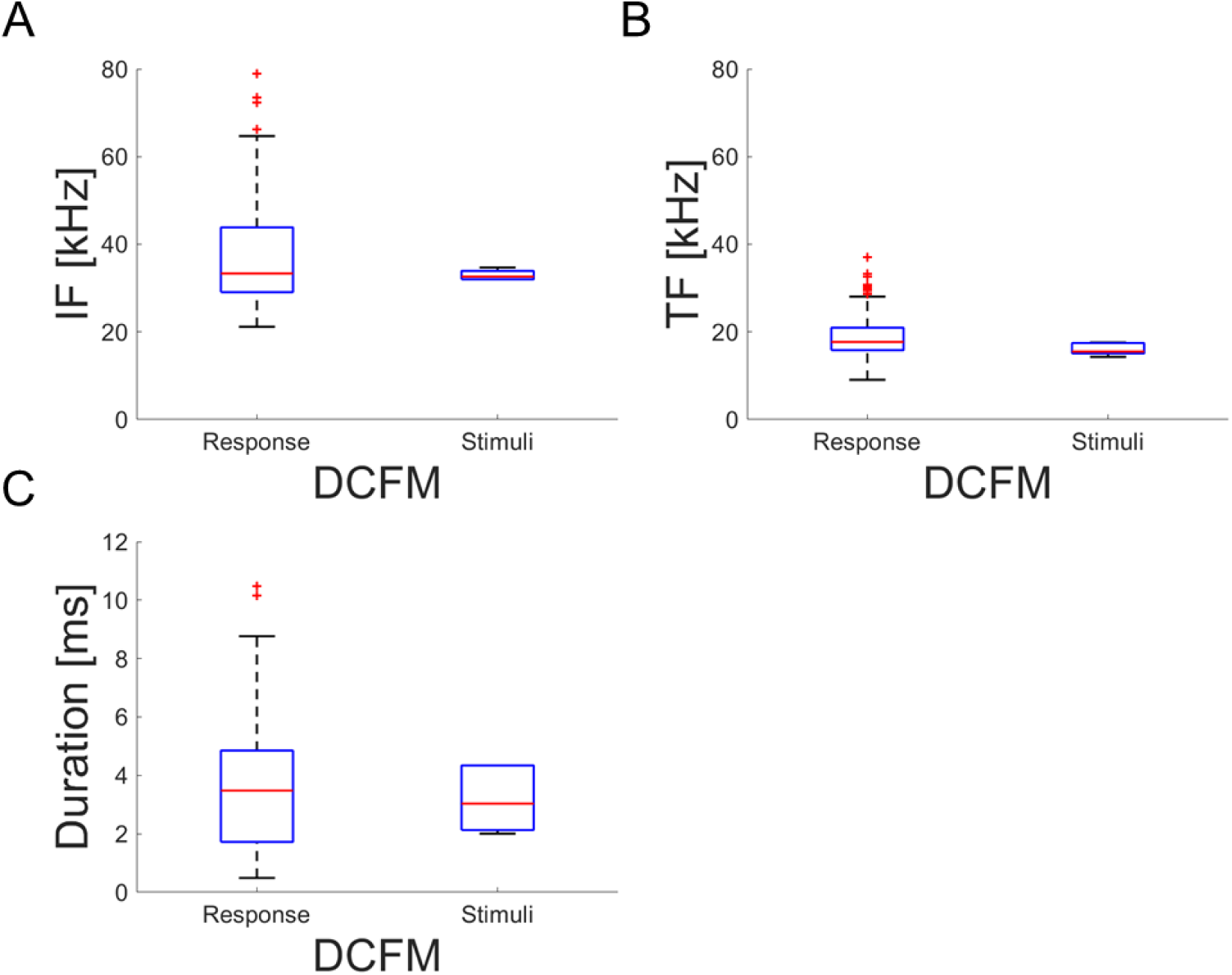
Comparison of acoustic parameters between the DCFM vocalizations of the donor bats in experiment I and the DC stimuli in experiment III. (A) the initial frequency, (B) the terminal frequency, and (C) the duration. Each box plot shows the median (red line), quartiles (the bottom and top edges of the boxes), and the whiskers (the minimum and maximum) (the top and the bottom tips of the vertical lines). The red cross marks indicate outliers (outside of the whiskers).

**Table S1.** Information from experiment I. Each trial includes all conditions, except for trial #11, which includes only a control condition. In the DB condition, in addition to the classification of behavioral responses and the reaction time (RT), the number of vocalization calls and the call ratio of each type are listed. Note that the RT is not listed when the response was “stay”.

| Trial # | Donor | Subject | DB response | DB RT [s] | # of calls | Ratio of EC [%] | Ratio of DCFM [%] | Ratio of DCNB [%] | Ratio of DCO [%] | NDB response | NDB RT [s] | Control response | Control RT [s] |
| --- | --- | --- | --- | --- | --- | --- | --- | --- | --- | --- | --- | --- | --- |
| 1 | M1 | F1 | Crawl | 81 | 1477 | 4 | 19 | 55 | 18 | Crawl | 159 | Crawl | 168 |
| 2 | M1 | F2 | Crawl | 165 | 1138 | 6 | 32 | 38 | 13 | Stay | - | Stay | - |
| 3 | M2 | F3 | Fly | 108 | 971 | 16 | 1 | 59 | 9 | Fly | 114 | Fly | 78 |
| 4 | M2 | F3 | Stay | - | 1190 | 0 | 8 | 72 | 8 | Fly | 114 | Fly | 33 |
| 5 | M2 | F4 | Stay | - | 729 | 13 | 3 | 60 | 8 | Stay | - | Crawl | 33 |
| 6 | M1 | F4 | Stay | - | 1532 | 6 | 30 | 50 | 13 | Fly | 51 | Stay | - |
| 7 | M2 | F4 | Stay | - | 1208 | 1 | 21 | 62 | 10 | Fly | 162 | Fly | 60 |
| 8 | M2 | M1 | Crawl | 27 | 1041 | 3 | 10 | 75 | 2 | Fly | 12 | Crawl | 18 |
| 9 | M2 | M1 | Stay | - | 1216 | 1 | 14 | 72 | 5 | Crawl | 69 | Crawl | 39 |
| 10 | M3 | M4 | Stay | - | 685 | 24 | 14 | 47 | 5 | Stay | - | Fly | 84 |
| 11 | - | M3 | - | - | - | - | - | - | - | - | - | Stay | - |

**Table S2.** Recording information details of experiment II. All pairs were of the same sex to avoid sexual interaction, and four of the pairs were kept in separate rearing cages from the time of capture for the subject bats. One subject-donor pair (M6 and M7) was kept in the same cage from the time of capture. Note that the M6 and M7 pair were tested twice because test #3 was terminated without stimulation of the control II.

| Test # | Donor | Subject | Cage | Recording time [s] | Included trials (Total) | DB | NDB | Control I | Control II |
| --- | --- | --- | --- | --- | --- | --- | --- | --- | --- |
| 1 | M4 | M5 | Separate | 1238 | 13 | 3 | 4 | 2 | 4 |
| 2 | M4 | M2 | Separate | 1196 | 9 | 2 | 3 | 2 | 2 |
| 3 | M6 | M7 | Same | 670 | 6 | 2 | 3 | 1 | 0 |
| 4 | M6 | M7 | Same | 646 | 6 | 2 | 2 | 0 | 2 |
| 5 | F5 | F7 | Separate | 849 | 7 | 2 | 3 | 1 | 1 |
| 6 | F6 | F7 | Separate | 1026 | 13 | 3 | 4 | 2 | 4 |
| 7 | F7 | F8 | Separate | 501 | 6 | 1 | 2 | 1 | 2 |

## Notes

### Competing Interest Statement

The authors have declared no competing interest.

